# Resolving structural diversity of Carbapenemase-producing gram-negative bacteria using single molecule sequencing

**DOI:** 10.1101/456897

**Authors:** Nicholas Noll, Eric Urich, Daniel Wüthrich, Vladimira Hinic, Adrian Egli, Richard A. Neher

**Affiliations:** Biozentrum, University of Basel, Basel, Switzerland; Swiss Institute of Bioinformatics, Basel, Switzerland; Division of Clinical Microbiology, University Hospital Basel, Basel, Switzerland; Applied Microbiology Research, Department of Biomedicine, University of Basel, Basel, Switzerland

## Abstract

Carbapenemase-producing bacteria are resistant against almost all commonly used betalactam and cephalosporin antibiotics and represent a growing public health crisis. Carbapenemases reside predominantly in mobile genetic elements and rapidly spread across genetic backgrounds and species boundaries. Here, we report more than one hundred finished, high quality genomes of carbapenemase producing *enterobacteriaceae, P. aeruginosa* and *A. baumannii* sequenced with Oxford Nanopore and Illumina technologies. We developed a number of high-throughput criteria to assess the quality of fully assembled genomes for which curated references do not exist. Using this diverse collection of closed genomes and plasmids, we demonstrate rapid movement of carbapenemase between genomic neighborhoods, sequence types, and across species boundaries with distinct patterns for different carbapenemases. Lastly, we present evidence of multiple ancestral recombination events between different *Enterobacteriaceae* MLSTs. Taken together, our samples suggest a hierarchical picture of genomic variation produced by the evolution of carbapenemase producing bacteria that will require new models to adequately understand and track.

---

The rapid global increase of multidrug-resistant organisms presents a major global health threat that will dramatically reduce the efficacy of antibiotics and thus constrain the number of effective treatments available to patients (Hawken and Snitkin, 2018; Logan and Weinstein, 2017). Routine surgergy or immunosuppression now carries the risk of untreatable life threating infections.

The global trend is acutely exemplified by the recent proliferation of microbial resistance to the carbapenems, a class of *β*-lactam antibiotics used for empiric treatement of infections with suspected multi-drug resistant bacteria and thus is of particular clinical importance. Carbapenemases, enzymes which hydrolyze carbapenems, have spread remarkably fast (Logan and Weinstein, 2017); the first *Klebsiella pneumoniae* carbapenemase (KPC) was isolated in the United States in 1996 and has since become the most common carbapenemase (DeLeo *et al.,* 2014; Holt *et al.,* 2015). A large variety of carbapenemases, e.g. NDM, OXA-48, IMP and VIM, have similarly risen in global frequency over this time-span due to the increase in usage of carbapenems in health care (Doi and Paterson, 2015; Logan and Weinstein, 2017; Nordmann *et al.,* 2011).

The spread of carbapenemases is putatively driven by both clonal expansion of specific strains, as well as horizontal transfer of mobile genetic elements within and across species boundaries (van Duin and Doi, 2017). It is commonly thought that the global spread of KPC is due to the dissemination of *K. pneumoniae* ST258, defined by multi-locus sequencing typing (MLST) (Munoz-Price *et al.,* 2013). However, studies have also found rapid KPC nosocomial outbreak dynamics originate primarily from prevalent plasmid and transposon transfer (Sheppard *et al.,* 2016). This pattern is not specific to KPC: the dissemination of OXA, TEM and NDM is equivalently accelerated in the short-term by their respective association to specific transposons that readily mobilize to different plasmid backbones (Holt *et al.,* 2015). Reconstruction of the evolutionary history of the outbreak is further complicated by an unknown rate of homologous recombination, shown to be extensive in the short term history of *Acinetobacter baumannii* as well as the ancestral history of current circulating *K. pneumoniae* strains (Chen *et al.,* 2014; Snitkin *et al.,* 2011).

Quantitative characterization of the global epidemiology of carbapenemase producers will require elucidation of the roles of horizontal transfer, clonal expansion, and homologous recombination in shaping the standing genetic variation in pathogenic bacteria (Holt *et al.,* 2015). While next generation sequencing has greatly increased our ability to resolve the short time dynamics of pathogen evolution, standard technologies fall short in one crucial aspect: the short reads generated by most sequencing technologies (e.g. Illumina), generically fail to assemble into complete genomes. Instead, genome assemblies are often fragmented into hundreds of contigs with breakpoints corresponding to repetitive regions such as mobile elements, a particularly severe limitation for the study of the molecular evolution of carbapenemase as they are usually associated with transposons. As a result, standard WGS in typically uninformative of the overall genomic neighborhood of each resistance gene and its evolutionary history.

Until recently, the production of fully assembled genomes required slow, expensive and labor intensive methods (Loman and Pallen, 2015). As a result, the number of high quality reference carbapenemase producing genomes is still small; for example, as of October 2018 the NCBI pathogen database only includes 112 assembled genomes (<5 contigs) of *K. pneumoniae* that contain a KPC *β*-lactamase. With the advent of high-throughput long read sequencing technology (PacBio, Oxford Nanopore) it has become possible to fully assemble bacterial genomes in a cost-effective, automated manner (Wick *et al.,* 2017a). Here we report and analyze the hybrid de-novo assemblies of 110 carbapenemase containing isolates of *Klebsiella pneumoniae, Escherichia coli, Acinetobacter baumannii,* and *Pseudomonas aeruginosa* isolated at the University Hospital Basel over the course of 8 years. Using a combination of Oxford Nanopore and Illumina MiSeq sequence data, we fully assembled 103 of the clinical isolates.

As traditional assembly metrics, e.g. N50, lack resolution for assessing the quality of fully assembled genomes, here we present diagnostics to assess both nucleotide and structural accuracy. A median 99.999% of each assembled genome showed no-evidence of gross sequencing error or misassembly, with the remainder likely due to the ambiguous mapping of short reads during assembly polishing.

Our large collection of high quality, complete assemblies provides a point of entry to comparatively study the genome evolution of antibiotic resistance to carbapenems. We show that the genomic contexts of the globally circulating carbapenemase genes exhibit a nested structure; we find both extensive inter-strain plasmid sharing as well as small-scale transpositions of the mobile elements flanking each carbapenemase between plasmids. Additionally, we find that a large fraction of the *Enterobacteriaceae* genomes are ancestral mosaics. This complex dynamics underscores the need for long-read sequencing in surveillance of resistance determinants in bacteria.

## MATERIALS AND METHODS

### Collection of bacterial isolates

The bacterial isolates originate from the culture collection of the Division of Clinical Microbiology at the University Hospital Basel. 110 well characterized carbapenem-resistant clinical isolates recovered from 2010 to 2017 within Basel, Switzerland were included in this study. A total of 87 *enterobacteriaceae,* 13 *P. aeruginosa* and 10 *A. baumannii.* The study has been approved by the local IRB (EKNZ Nr 2017-00222).

Strains were either detected in screening from colonized patients (Hinić *et al.,* 2017, 2018) or in routine diagnostics from an infection most often using VITEK 2 (bioMérieux). All carbapenemase-producing strains were confirmed by either in house PCR (Poirel *et al.,* 2011; Woodford *et al.,* 2006), Xpert-Carba™ (GeneXpert, Cepheid) or eazyplex™ superbug CRE (Amplex Biosystems). Both matrix assisted laser desorption ionization-time of flight (MALDI-TOF) mass spectrometry on a Bruker microflex system (Bruker Daltonics, Bremen) and routine antimicrobial susceptibility testing using VITEK™ was used for species identification

### DNA extraction

Strains were stored at microbial storage tube Microbank™ (PRO-LAB Diagnostics, US) at −80°C, re-cultured on Columbia agar with 5% sheep blood (bioMérieux, France) and incubated at 37°C under atmospheric conditions with 5% CO_2_ for 18-24 hours. A filled inoculation loop of colonies were collected and suspended in 2mL Phosphate-Buffered Saline (Sigma-Aldrich Chemie GmbH, Switzerland). This solution was centrifuged for 10 minutes at 13’000 rpm to collect the bacterial pellet.

**TABLE I.**
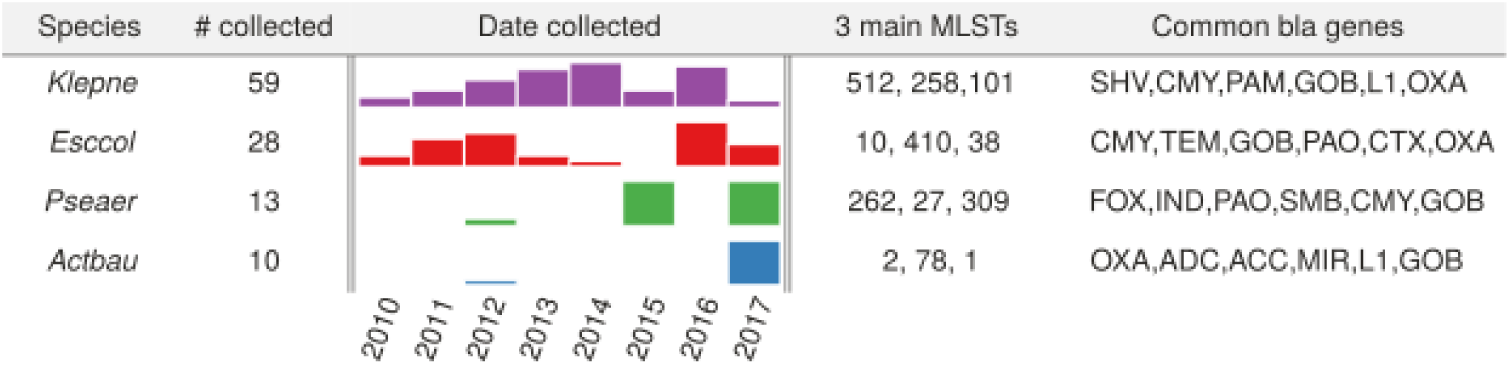
Sample overview. Species breakdown, collection dates, main sequence types and *β*-lactamases of the samples sequenced in this study.

The DNA extraction was performed from a bacterial pellet using the QIAamp^®^ DNA mini kit (QIAGEN) on the QIAcube^®^ robot (QIAGEN) according to the manufacturer’s protocol. The bacterial DNA was eluted in 150*µ*l AE buffer and stored at −20°C. The DNA concentration was measured using Qubit^®^ 3.0 fluorometer (Invitrogen, US) with the Qubit™ dsDNA HS assay kit.

### Long-read sequencing

Each library was prepared using a custom protocol based upon the ONT 1D ligation sequencing kit (LSK-108) supplemented with the native barcoding expansion kit (EXP-NBD103). The optional shearing and repair step of the protocol was omitted to obtain longer reads. Since adapter ligations during library preparation and nanopore sequencing are foremost sensitive to the molarity of sequencing library, we increased the amount of total DNA material used. Each isolates’ extracted DNA was diluted to 1*µ*g/*µ*L in 50*µ*L nuclease free water (NFW). 7*µ*L of NEBnext Ultra II End-repair/dA-tailing buffer and 3*µ*L of NEBnext Ultra II End-repair/dA-tailing enzyme mix were added to each sample and incubated at 20°C for 5 min and 65°C for 5 min. 60*µ*L of AM-Pure XP beads were added to each sample and then each DNA sample was washed using 70% EtOH and eluted into 25*µ*L of NFW. 2.5*µ*L of each barcode plus 25*µ*L of Blunt/TA ligase master mix was added to each sample and incubated at room temperature for 10 minutes. The samples were immediately pooled together and then washed by adding 250*µ*L of AM-Pure XP beads to the pooled sample. Following another wash with 70% EtOH, all DNA was eluted into 51 *µ*L. 1*µ*L was used to measure DNA concentration with a Qubit fluorometer. The final sample was diluted to a DNA concentration of 35ng/*µ*L. Adapter ligation was prepared by adding 20*µ*L BAM, 30*µ*L Ultra II ligation master mix, and 1*µ*L ligation enhancer to the pooled sample. The library was spun down and incubated at 10 minutes. The remaining steps followed the LSK-108 kit protocol.

All 110 samples were sequenced over 15 different runs and averaged ~ 9 samples/flowcell. 25 samples had to be re-sequenced due to low coverage resulting in poor assembly. All resulting fast5 read files were base-called using ONT’s albacore command line tool (v2.0.2). Albacore was run using barcode demultiplexing with Fastq output. Porechop (v 0.2.3) was used on the directory output from Albacore and only reads that both Albacore and porechop agreed upon were kept and stored in one Fastq file to be used for subsequent assembly.

The demultiplexed and adapter-trimmed sequencing reads are available at ENA under bioproject number PRJEB28660.

### Short-read sequencing

The DNA from cultured isolates was extracted using the EZ1 DNA tissue kit on an EZ1 Advanced XL robotic system (Qiagen). The extracted DNA was processed using the Nextera XT library preparation kit (Illumina). The resulting library was sequenced using a MiSeq Illumina platform (accredited with ISO/IEC norm 17025) with 2×300 paired-end sequencing as previously described (Piso *et al.,* 2017)

The demultiplexed sequencing reads are available at ENA under Bioproject number PRJEB28660

### Genome assembly, annotation and sequence typing

Each ONT read set was assembled using Canu (v1.5) (Koren *et al.,* 2017) on a high-performance computing cluster (sciCORE at the University of Basel). If an isolate was sequenced multiple times, all available read sets were given to the assembler. The corresponding Illumina reads were mapped to the resultant assembly using bwa-mem for multiple rounds of polishing with Pilon (v1.22) (Walker *et al.,* 2014).

Additionally, each assembly was additionally performed using Unicycler (v0.4.2) (Wick *et al.,* 2017b). Assemblies between Unicycler and Canu were compared using the genome alignment mode of Minimap2 (Li, 2018). In general, Canu produced more contiguous assemblies than Unicycler, most likely because most the assemblies of the Illumina data contained many dead ends. This is likely due to the small insert sizes of our paired-end short read libraries, see S.I. for more detail. Comparison between Canu and Unicycler also provided a sanity check on assembly methods as in general both assemblies agreed 99.9% with each other at the nucleotide level. Conversely, Unicycler did assemble a few small (< 20 kb) plasmids that did not map to any corresponding plasmid in the Canu assembly, either because it was not present in our ONT library or because Canu excluded these reads. When this occurred, the Unicycler contig was appended to the Canu assembly.

All assembled genomes were annotated using prokka (v1.12) (Seemann, 2014) supplemented with a protein database of *β*-lactamases from ResFinder (Zankari *et al.,* 2012), as well as an HMM to annotate insertion sequences from ISEscan (Xie and Tang, 2017). Additionally all plasmids were typed using PlasmidFinder (Carattoli *et al.,* 2014). Lastly each isolate was typed using MLST (Jolley and Maiden, 2010; Seemann, 2018).

The annotated assemblies are available in Genbank under accession numbers ERZ777024- ERZ777123.

### Synteny Alignments

Gene synteny in the neighborhood of carbapenemase genes was assessed by representing plasmids or the 100 flanking genes (for chromosomal carbapenemase genes) as strings of orthologous gene clusters. These gene clusters were inferred using PanX (Ding *et al.,* 2018). PanX first performs an all-to-all alignment of proteins using DIAMOND (Buchfink *et al.,* 2015) and then performs the Markov Cluster Algorithm on the resulting graph of e-values. Paralogous clusters are delineated by cutting anomalously long branches in the resulting phylogenetic tree.

Each gene in the original sequences was then represented by an ID unique to its gene cluster. All pairs of the coarse-grained sequences were then aligned using the pairwise local alignment algorithm of Seqan (Döring *et al.,* 2008) exposed to python via the Cython interface (available on GitHub). For circular plasmids, we repeated the alignment for all possible rotations of the sequence and its reverse complement.

The resulting matrix of edit distances was clustered into an ultrametric tree, using the UPGMA algorithm, such that genomic contexts were hierarchical clusters of synteny/genomic environments based upon ‘edit’ distance of the strings made of orthologous genes.

### Core genome trees and homologous recombination analysis

The core genome for each species of collected isolates were built using the PanX pipeline using default parameters(Ding *et al.,* 2018); the core genome tree was subsequently inferred using FastTree (v2.1.9) (Price *et al.,* 2010), using the ‘-gtr’ and ‘-gamma’ flags, from the concatenated nucleotide alignment of all core genes. The resulting branch lengths of all phylogenetic trees were further refined using TreeTime (Sagulenko *et al.,* 2018). All tree distances obtained from this process were used as a proxy for evolutionary distance in the subsequent analyses.

The core tree allowed inference of mutational events; mutations were estimated using message passing in a maximum joint likelihood framework using the TreeAnc class of TreeTime (Sagulenko *et al.,* 2018) Mutations that were observed more than once within the tree were classified as homoplasies. Both homoplasic density and total SNP density were obtained by averaging the number of each respective event over the concatenated core genes in a sliding window of size 5 kb with a custom Python script.

A hash map was defined - i.e. each orthologous group obtained from PanX was given a unique integer identifier - this allowed for quick comparison between the gene content of different isolates. Additionally, the defined hash corresponding to the core genes were used as markers to identify global rearrangements between pairs of isolates. Broken relative ordering of the defined hash was inferred as a rearrangement event. Specifically, all isolates were compared to a defined reference, chosen from the dominant subtype for each species: the internal numbered isolate 13, 70, 116, and 47 were chosen for *E. coli, K. pneumoniae, P. aeruginosa,* and *A. baumannii* respectively. Continuous deviations of the gene order, defined relative to the reference, for the remaining isolates were utilized as a proxy for a single rearrangement event.

To supplement the above homoplasy analysis, a tree scan was performed on each core genome alignment. Each alignment was partitioned into approximately 10 kb blocks, taking care to not extent blocks past rearrangement events as detected above. The resulting sub-alignments were fed into FastTree (Price *et al.,* 2010) with the same parameters as used for the full core genome tree to build local trees. All local trees were compared to each other using the Robinson-Foulds metric, implemented using the Dendropy (v4.2.0) module (Sukumaran and Holder, 2010) The resulting tree distance matrix was clustered using the UPGMA algorithm implemented in Scipy (v1.0.1) (Oliphant, 2007). Flat clusters were obtained by looking for clusters greater than 25% of the maximum RF distance away. All 10 kb blocks associated with one flat cluster were concatenated and passed to FastTree to better resolve the tree corresponding to each partition.

## RESULTS

### Assembly of complete genomes

Despite the fact that long reads have facilitated the automation of complete bacterial genomes (Wick *et al.,* 2017a), genomic assembly still has many potential pitfalls (Schmid *et al.,* 2018). As such, stringent quality metrics are necessary; as the price of whole genome sequencing continues to decline, the number of assemblies per project will surpass what is feasible for manual inspection and validation. Traditional assembly metrics, e.g. N50 and L50, assess solely the fragmentation of the final assembly. As most of long read assemblies of bacterial genomes readily assemble into circular chromosomes, these metrics were uninformative for our present purposes. Instead, we utilized metrics that directly quantify the *accuracy* or *quality* of the assembly relative to three different sources of potential error: (i) single base errors in the nucleotide sequence, (ii) small indel errors (<15bp), and (iii) errors in the global structure of the assembly, e.g. collapsed repeats. Importantly, these metrics should quantity accuracy without reference to the correct sequence which is unknown *a priori.*

#### Substitution errors

Accuracy of individual bases in the assembled sequence primarily depends on the accuracy of the sequencing technology and the degree to which these errors are random as opposed to systematic. Such errors are of particular concern for genomes assembled from solely noisy long reads (Lo-man *et al.,* 2015); we estimated the nucleotide error rate to be ~ 13.6% from the ONT data of all our bacterial genomes (see Fig. 1 B), consistent with previously reported numbers (Rang *et al.,* 2018). Importantly, the majority of the measured errors are attributed to false insertions or deletions relative to polished assembly; false nucleotide substitutions were found to occur only at rates of about 3%. In contrast, the indel error rate of the Illumina technology has been measured to about 10^-5^ (Zanini *et al.,* 2017). ONT errors were found to be strongly correlated in space and enriched within homopolymeric stretches of the genome, also reported in (Loman *et al.,* 2015). The consensus taken over a deep coverage of long reads polished away many of these errors, however, as previously observed (Mikheyev and Tin, 2014), nanopore-exclusive assemblies were found to have an enriched rate of indel errors that dramatically affected downstream ORF discovery and subsequent annotation. Thus, at present, the only way to guarantee accurate assemblies is to perform a hybrid assembly using noisy long read for long-range scaffolding information supplemented with Illumina data for short read accuracy. We refer the interested reader to the S.I. for a more in-depth discussion of sequencing errors measured in the raw reads of both technologies reads and different assemblies.

**FIG. 1.**
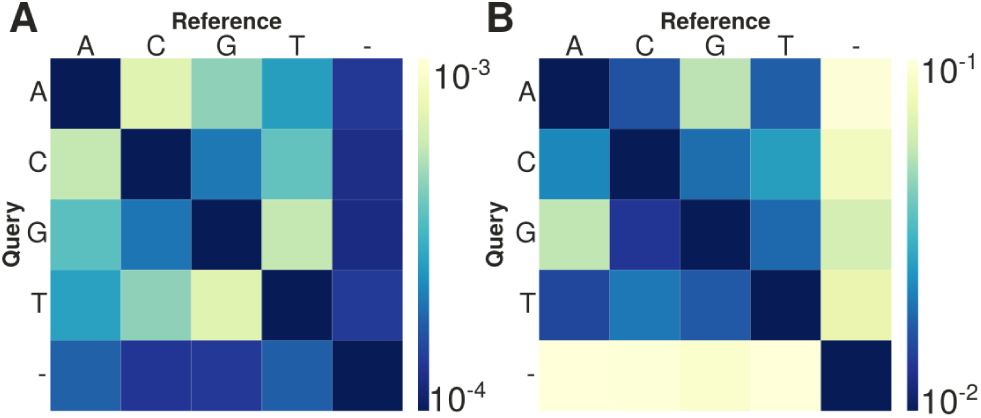
Sequencing error rates. Error rates of raw reads for A) Illumina and B) ONT reads, measured relative to the final polished assembly, averaged over all clinical isolates (note the 100-fold difference in the color scale of panels A & B). Errors in ONT reads are dominated by false indels, while such errors are very rare in Illumina data.

In order to quantify substitution errors without knowing the true sequence, we compared diversity in the pile-up of Illumina short reads with the expectation from the known Illumina error profile. For simplicity, we ignore the existent and comparatively small context biases of the Illumina platform and assumed a constant Poisson rate of errors. Specifically, as each final genome assembly was polished under multiple rounds of Pilon, each base-pair should have an error rate consistent with Illumina technology, estimated to be *p* ~ .12% from Fig. 1A. If, however, polishing failed due to ambiguous short read mapping, collapsed regions within the assembly, or a region with anomalously high error rate preventing short read mapping, then some columns within the pile-up will be more diverse than expected by the error model. This null hypothesis was quantitatively tested by computing the binomial likelihood of observing *n_i_* Illumina reads that differ from the assembly in alignment columns *i,* conditional on the observed coverage *c_i_,* on the polished genome assembly.

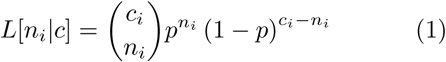

The empirically measured coverage distribution was used in combination with Bayes’ rule to compute likelihood for each column *i.* The likelihoods of all columns were then binned into a histogram; in absence of errors in the assembly, the fraction of sites within each bin should be equal to the likelihood. In general, we find the expected relationship between likelihood and fraction of sites over 6 orders of magnitude, see Fig. 2A; however we do observe a consistent enrichment of sites at likelihoods lower than 10^-6^ in most of our assemblies. Specifically, as shown in the inset of Fig. 2A, approximately 5.10^-5^ of all pile-up columns have anomalously low likelihoods inconsistent with the sequencing error rate. These sites were clustered in specific regions of each genome and tended to fall into repetitive regions where mapping and thus assembly polishing remains ambiguous.

**FIG. 2.**
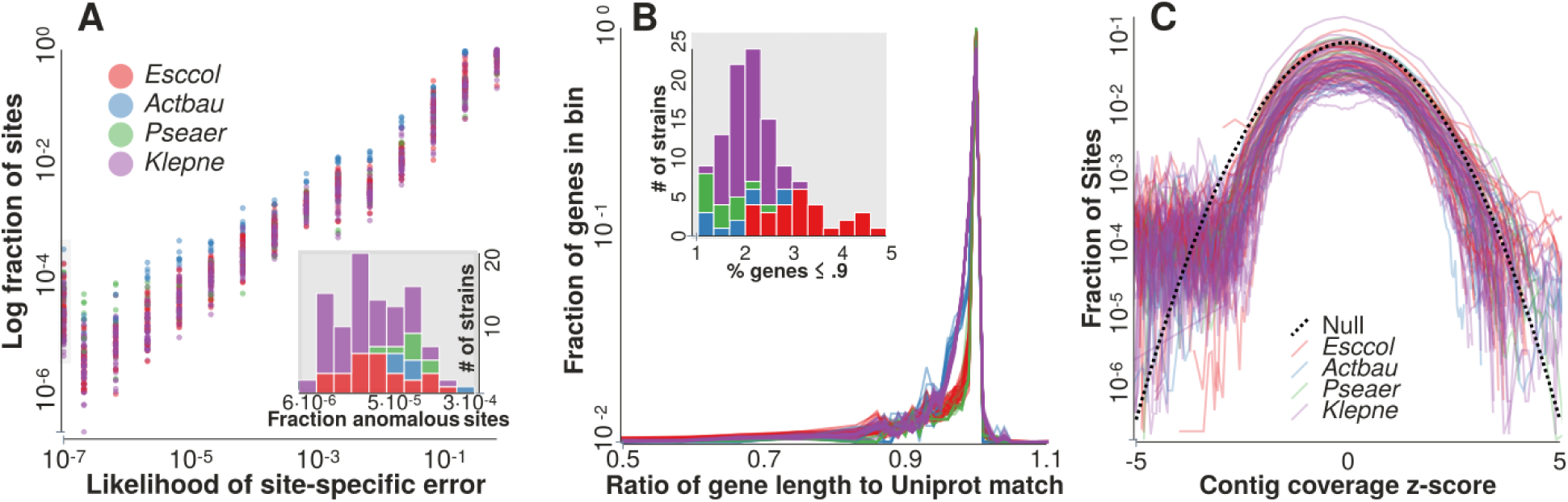
Assembly statistics and quality. A) Diversity in the pile-up of Illumina short reads mapped to the hybrid assembly is consistent with a short read error rate of 0.12%. A perfect fit to the null expectation would show as a straight line in this plot - the main deviation can be found in the pileup at the left. Around 3 in 10^5^ pile-up columns show significantly higher diversity than expected, exhibited in the enrichment of unlikely sites. Such columns tend to fall within repetitive regions where mapping and assembly polishing remains ambiguous. B) The great majority of annotated proteins have a hit of equal length in SwissPROT, suggesting frameshift errors are rare within our assemblies. Conservatively, assuming no pseudogenization we estimate about .2% of genes are prematurely truncated. Note that the y-scale is logarithmic, greatly emphasizing the rare truncated proteins. The inset shows distribution of annotated proteins that are shorter than 90% of their closest SwissPROT hit. C) Long read coverage follows the expected Poisson distribution over 3 standard deviations. The graph 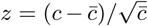 where 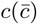 are (average) coverage.

#### Short indels and gene prediction

Many sequencing technologies, including ONT, PacBio, 454 and Ion-Torrent, struggle with homopolymeric tracts resulting in indel-errors in such regions. Such errors are not easily corrected by consensus building of many reads since there are systematic biases, either in sequencing or consensus building. It should be expected that assemblies using only such technologies will suffer from frequent short indel errors, confounding gene prediction via premature stop codon and reading frame shifts. Illumina sequencers, in contrast, have a per base indel rate below 10^-5^, see Fig. 1A, thus polishing a long-read assembly with short reads is expected to remove most indel errors.

We pursued a quantitative estimate of the effect of the remaining indels errors on our final genome assemblies. Specifically, indel errors are known to dramatically affect downstream automated gene prediction and annotation steps by artificially producing a frame-shifts or premature stop codons within the protein coding regions. In order to get a quantitative handle on the occurrence rate of such errors, we aligned all proteins for each annotated genome to the manually curated SwissPROT database (The UniProt Consortium, 2017) using DIAMOND (Buchfink *et al.,* 2015) and compared the length of our predicted protein to the top match in the database. The resulting distribution of ratios is shown in Fig. 2C. Following (Watson, 2018), this distribution was used to estimate the fraction of prematurely truncated genes within our assemblies. To accomplish this, we analyzed the fraction of predicted genes that differed by more than 10% from the top match in SwissPROT, shown within the inset of Fig. 2C. We estimate that approximately ~ 2% of genes within each genome are prematurely shortened, generally consistent with expected pseudo-gene content (Liu *et al.,* 2004).

#### Large scale assembly accuracy

Lastly, global assembly accuracy was quantified by using fluctuations from the null distribution of ONT read coverage. Specifically, it has been observed that nanopore uniformly samples the sequenced genome and that coverage does not fluctuate widely as it does for Illumina Nextera XT (e.g. with GC content) (Loman *et al.,* 2015). We confirmed this across the 110 sequenced samples from different species, as shown in Fig. 2C. The figure shows histogram of deviations from the average coverage for each contig across all samples, rescaled by the standard deviation expected from perfect Poisson sampling. Global mis-assemblies and collapsed regions should show up as outliers in the coverage distribution. For about three standard deviations, this histogram follows the null expectation from randomly sampling at constant density suggesting that most of the assembly has a one-to-one correspondence to the genome of the isolate.

**TABLE II.**
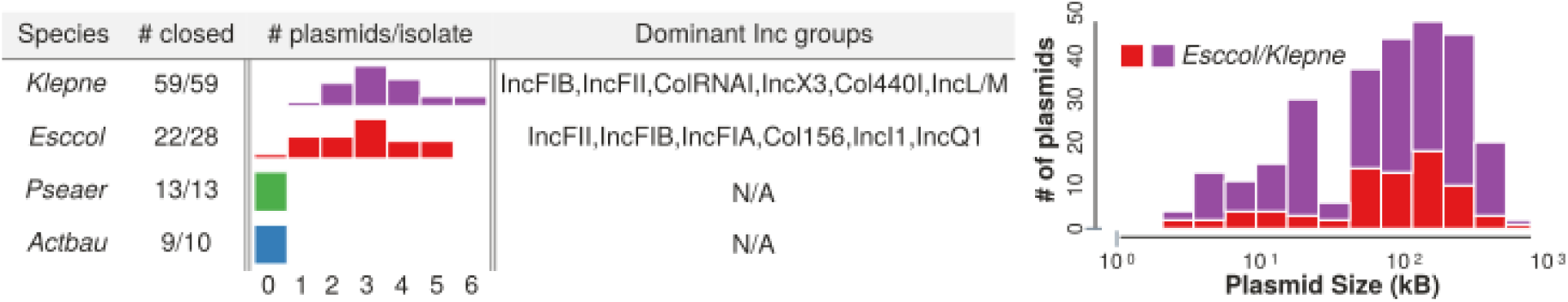
Chromosome and plasmid assembly characterization. A) Table containing information about assembly completion and number of found plasmids per individual. Interestingly, no plasmids were found in either the *A. baumannii* or *P. aeruginosa* isolates. In contrast, most *E. coli* and *K. pneumoniae* isolates contained between 1 and 5 plasmids. B) The length distribution of plasmids found in the sample. We observe two distinct size distributions: large (~ 100 kb) and small (~ 200*kb*), consistent with prior observations.

However, there is a systematic low-coverage tail in Fig. 2C where the distribution that deviates from the expectation. This tail, on average, represents approximately 5 · 10^−5^ of the entire genome.

In summary, the most prevalent errors are due to imperfect polishing and indel-correction, which fails in regions where short read coverage is anomalously low (for example in very AT rich regions), in repetitive regions were mapping is ambiguous, and in regions of where the ‘ONT-only assembly’ has too many errors for standard mappers. While such regions likely exist in most of our genomes, our analyses here show that these problems are restricted to a small number of sites. This number is consistent with that obtained via our short read data. Outside of roughly 200 – 800 nucleotides per genome, our final assemblies are internally consistent with the raw data.

### Carbapenemases reside in diverse set of genomic backgrounds

It is largely thought that short term dissemination of carbapenemases, and *β*-lactamases in general, is driven by the clonal dissemination of resistant lineages such that transmission chains involve a single pathogenic lineage (Lu *et al.,* 2018; Munoz-Price *et al.,* 2013; Pournaras *et al.,* 2009) Importantly, this has influenced surveillance strategies to focus on tracking strains via ‘sequence types’ using techniques such as MLST and pulsed-field gel electrophoresis which are gradually replaced by WGS and core genome MLST (Hawken and Snitkin, 2018).

However, it is becoming increasingly clear that carbapenemases undergo rapid horizontal transfer due to their association to mobile genetic elements. Most carbapenemases (and resistance genes in general) are localized on conjugative plasmids that can be easily shared amongst bacteria. Additionally, carbapenemases are typically integrated within transposable elements that allow for rapid mobilization to other regions in the containing genome (Sheppard *et al.,* 2016). If either the rate of plasmid transfer or transposition is comparable to time-scale on which a clonal lineages spreads, then the outbreak dynamics is no longer well-documented by solely tracking sequence types. Rather, one would need to track the resistance determinant along with its genomic background (Wang *et al.,* 2018). Hence determining the ‘correct’ molecular unit of resistance that researchers should survey is of critical importance necessitating long-read sequencing.

Even short read WGS is ill-suited for linking resistance genes to their genetic background due to limitations of de-novo assembly of highly-repetitive regions. As shown in Fig. 3, the median contig length of a resistance gene in an Illumina-only assembly is roughly equal to the 10^th^ percentile of contig length obtained by our hybrid assembly. Lastly, reference-based techniques that utilize short reads are inadequate as such analyses *assume* relatively stable genomic structure of each plasmid background.

**FIG. 3.**
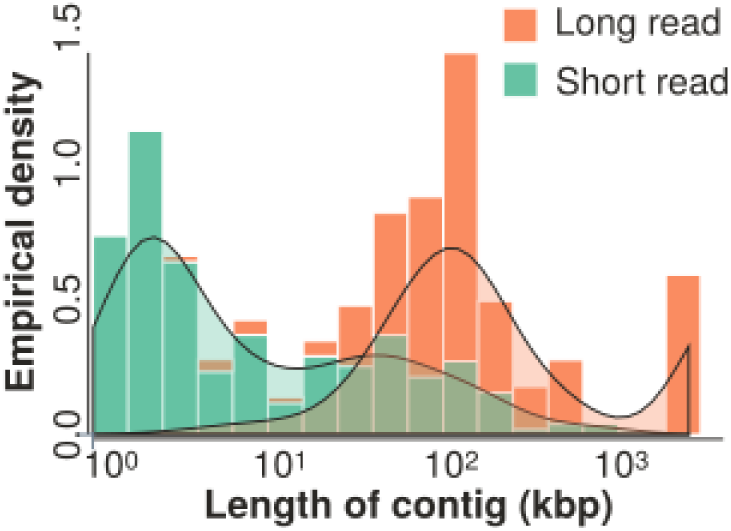
Length of contigs containing either carbapenemase and ESBL genes. Stacked histograms of the length of contig containing all genes annotated as a Carbapenemase or ESBL for a short read and hybrid assembly pipeline, shown in green and orange respectively. All plasmid encoded genes are fragmented into contigs of roughly 4 genes in the short read assembly, while fully resolved using ONT. Smoothed densities of the two distributions are shown in black.

High-quality assemblies allow us to investigate the relative importance of different mechanisms by which carbapenem resistance spreads. As shown in Table. II, 103/110 genomes were fully assembled - i.e. the largest contig was circular and had a size compatible with typical genome size of the organism. For all completed assemblies, the remaining contigs were assumed to be putative plasmids and verified against the PlasmidFinder database (Carattoli *et al.,* 2014). The distribution of plasmid number per isolate can be found in table. II, along with the most frequent incompatibility groups for each species. The table of all resulting incompatibility types for each isolate can be found in the S.I.

Evolutionary relationships between present-day isolates are typically reconstructed by similarity within alignments of homologous sequences, and only utilize single nucleotide substitutions and small indels. In the case of the carbapenemase pandemic, however, the genetic context evolves by transposition such that identical gene sequences are found in very different genomic neighborhoods. The genes flanking the resistance determinant and their order is therefore a more sensitive measure of evolutionary distance.

Given the large number of assembled isolates, we designed a high-throughput pipeline able to assess the similarity of carbapenemase genetic contexts. Instead of aligning the nucleotide sequences directly, we clustered all protein coding regions on the plasmid of each carbapenemase gene (or the flanking 100 genes if it was found integrated in the chromosome) into orthologous clusters using the PanX pipeline (Ding *et al.,* 2018). Each plasmid was coarse-grained a string of orthologous clusters, where each defined gene is represented by the unique id of the corresponding gene cluster. These gene-order sequences were aligned pairwise and the resulting edit distances used to construct ultra-metric trees (see materials and methods for details). Critically, this allowed us to quantify the similarity between each genomic contexts that define long stretches of synteny, independent of whether the gene falls on a plasmid or chromosome, in a scalable, computationally efficient manner.

#### Most KPC is found on two related plasmids

The tree obtained for all plasmids containing *blaKPC* is shown in Fig. 4A. We note that length of the longest branch from root to a leaf within the structural tree represents an edit distance of approximately .9 – this reflects the fact that *blaKPC* was universally found within the *tn4401 transposon* (Sheppard *et al.,* 2016) for all isolates, and as such even distinct plasmids could be locally aligned in the vicinity of the gene of interest.

**FIG. 4.**
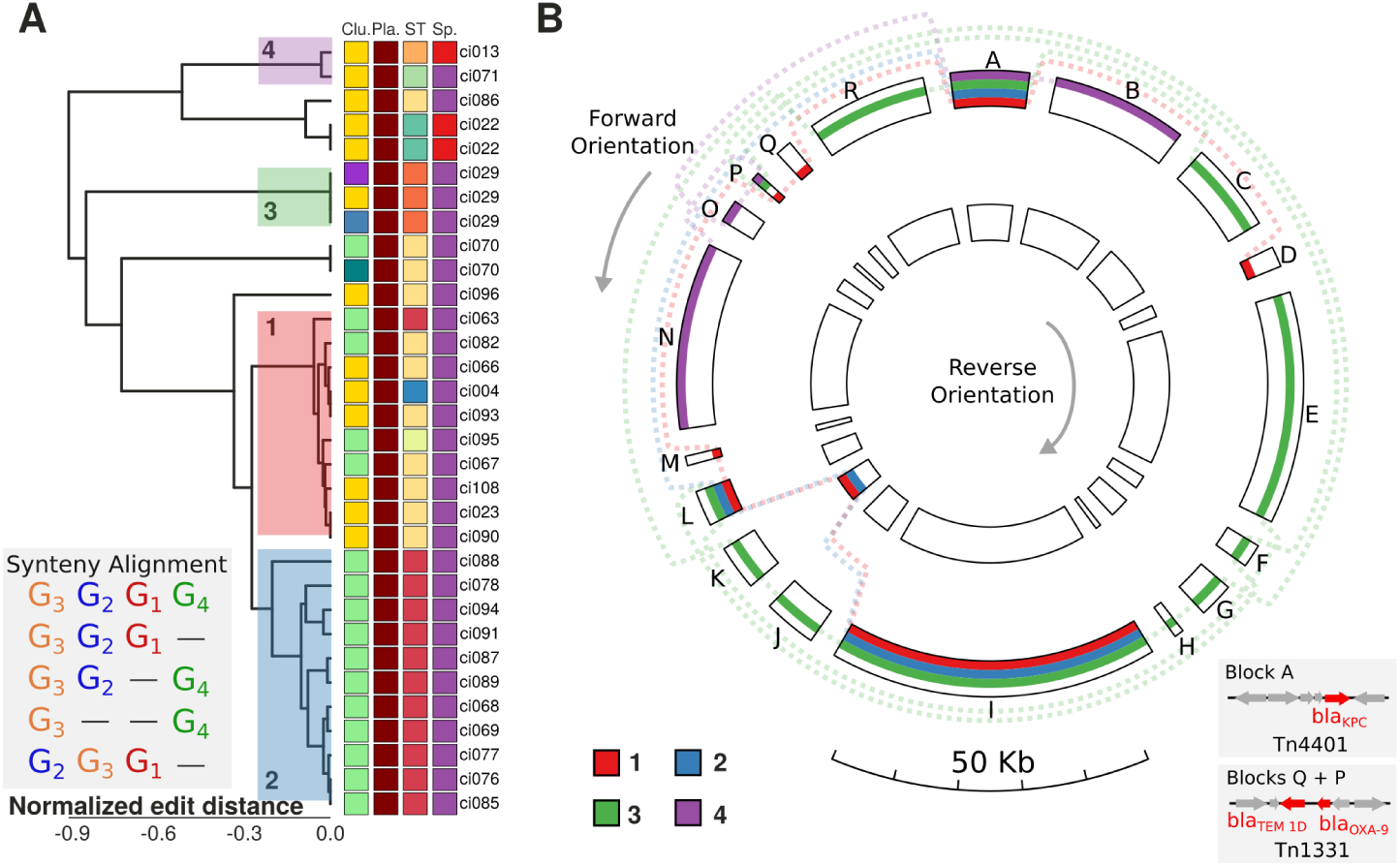
Genomic backgrounds of KPC. A) A dendrogram produced from the matrix of all pairwise edit distances obtained by local alignment of the gene alphabet as shown in the inset. We see 6 main structural clusters of plasmids from our 31 samples. To the left of the tree is shown a colored representation of meta data to compare to the dendrogram: genetic clusters (Clu.), if it was found on a plasmid (denoted Pla. where red denotes it was on plasmid, tan if within chromosome), the sequence type determined by MLST (ST), and the bacterial species (Sp.) (color code as in Tab. I) are shown left to right. B) A graphical depiction of a multiple genome alignment of four representative plasmids containing KPC as marked in A. Each color represents an plasmid chosen from the corresponding colored clade in A. Homologous stretches are denoted as lines occupying the same block, defined within the Mayday program. Block A corresponds to the Tn4401 transposon that all blaKPC genes were contained within in our sample. Additionally, block I represents the plasmid replication machinery and origin of transfer.

Roughly two thirds of all observed genomic contexts of *blaKPC* were found to be clustered within one of two closely ‘related’ plasmid backbones, denoted as clades 1 (red) and 2 (blue) in Fig. 4. Both clades predominately contained MLSTs 258 and 512, shown respectively in yellow and red in the third column of meta data to the right of the structure tree of Fig. 4A. Variation within each clade is predominately driven by small-scale point insertions of a few genes throughout the plasmid that result in gaps within the syntenic alignment. We note this is likely a combination of two factors - both real genetic presence absence variation between plasmids as well as issues related to ab initio gene prediction as discussed in the main section.

However the major structural difference between both clades is the insertion of an additional transposon carrying *blaTEM-1d* and *blaOXA-9* directly upstream of *tn4401* in the lower clade (highlighted in blue). We note that the plasmid associated with the structural clade 2 was found *only* within genomes classified as ST512 while clade 1 (highlighted in red) was predominately associated with ST258, with three other types - including ST512 - also found with the plasmid cluster.

The subtype diversity within a plasmid cluster suggests frequent plasmid exchange and plasticity, however the current data is too sparse to produce a reliable estimation of quantitative rates. Overall, clusters defined by the sequence of the KPC gene itself (the first column of the meta-data matrix in Fig. 4) are mostly inconsistent with clusters defined based on plasmid structure. The KPC sequences does, however, suggest that the ancestral plasmid of clade 2 resulted from the transposition of the *blaTEM-1d* transposon into one of plasmid variants from clade 1.

The remaining 11 KPC genes sit on 5 very distinct plasmids with few alignable regions outside of the *tn4401* transposon found universally within all isolates containing KPC, see Fig. 4. The isolate 29 (shown in green) and the two large clades discussed above, for example, differ greatly in gene content and structure as shown in Fig. 4B (created using GenomeRing (Herbig *et al.,* 2012)). Specifically, they are only homologous within the *tn4401* transposon (labeled block A in Fig. 4B) and the plasmid replication machinery (shown as block I). Additionally, the plasmid of isolate 29 also exhibited two large syntenic regions with structural clades 1 and 2, labeled blocks L and K in Fig. 4B), which was annotated as a transposon conferring mercuric resistance and block I, the origin of replication for each plasmid, in Fig. 4B. We note that isolate 29 has two duplications of *blaKPC* upstream of the *tn4401 transposon,* denoted as block R.

#### Extensive diversity of genomic backgrounds of NDM

While we found KPC exclusively on plasmids, six of the 16 NDM genes were found in the chromosome of *K. pneumoniae* or *E. coli* isolates (Fig. 5A). Additionally, seven NDM genes where found in essentially unique genomic backgrounds. The remaining 9 isolates fell into one cluster of three and one cluster of six sequences. The latter contains one NDM gene that is integrated into the chromosome, while the remaining five sequences are on plasmids with moderate structural align-ability (edit distance 0.1-0.3). As was found in KPC, all NDM genes were associated with a single transposon *tn125* which acts as the root of the dendrogram shown in Fig. 5A. This suggests a picture of rapid transposition of the NDM carbapenemase, in accordance with previous studies that have found a diverse set of NDM-encoding plasmids (Johnson and Woodford, 2013)

**FIG. 5.**
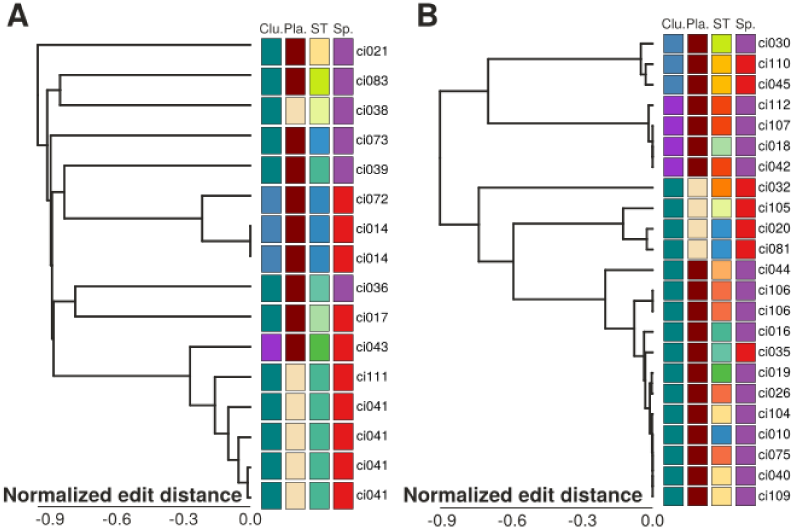
Genomic backgrounds of carbapenemase genes. A) NDM was found in a diverse set of genetic environments. Outside of the bottom structural clade, no regions were alignable outside of the NDM transposon. Additionally, we note that the bottom plasmid was found to be integrated within the genome. B) Conversely, OXA-48 has well defined structural clades that are tightly associated with phylogenetic clusters made from the gene alignment (first column). Interestingly, we see no strong association to ST, suggesting rampant plasmid transfer between individuals.

#### Rapid mixing of sequence type and genomic background of OXA-48

Fig. 5B shows the clustering of OXA-48 genes based on the edit distance obtained by the syntenic alignment of their flanking sequence. We find a very different picture relative to NDM; twelve out of twenty-four OXA-48 genes are part of a large group of structurally similar plasmids. However surprisingly, this tight cluster at the plasmid level contains six different sequence types of *K. pneumoniae* and one *E. coli* isolate. Additionally, it closest cousin clade within the dendrogram was found within the chromosome of an *E. coli* isolate, suggesting a partial transposition of the conjugative plasmid. The remaining OXA-48 genes fall into four groups (at edit distance 0.5) with one to four sequences each. Yet, even these small clusters contain multiple sequence types and species. This complete discordance of the similarity relationships of the neighborhood of OXA-48 and the similarity between the core genomes of the carrying strain underscores its extraordinary mobility. Discussion of ESBL backgrounds can be found in the SI.

### Characterization of the pan-genome

Traditionally, the global spread of bacterial pathogens is tracked using multi-locus sequences types, i.e. combinations of alleles of a number of informative core genes. Since such typing lacks resolution to track transmission chains of outbreaks, bacterial typing is rapidly shifting towards WGS via techniques such as cgMLST(Carleton and Gerner-Smidt, 2016). The relationship between isolates is reconstructed via a phylogenetic tree, obtained from the concatenation of the core genome, that putatively captures the clonal structure of the pathogen. The degree to which such trees provide an accurate, comprehensive summary of the evolution of strains is unclear. While the core genome tree likely provides an accurate representation of population structure and history over the time scale of an outbreak, homologous recombination, horizontal transfer, and genomic rearrangements will confound tree reconstruction of diverse strains.

#### Diversification of gene content and gene order

To assess the prevalence of gene gain/loss events in the context of our clinic isolates, we constructed the pan genome for each species using PanX (Ding *et al.,* 2018). The core genome sizes were found to be 3180, 3410, 2378, and 4411 genes for *E. coli, K. pneumoniae, A. baumannii,* and *P. aeruginosa* respectively, roughly 1/3 of all observed genes for *E. coli* and *K. pneumoniae* and 1/2 for *A. baumannii* and *P. aeruginosa.* This mostly reflects the relative genetic diversity and size of our samples.

We investigated the relationship between pairwise genetic divergence in core genes and the number of genes found in one isolate but not the other; the resulting scatter is shown in Fig. 6A. Interestingly, we find a sublinear relationship between genetic distance and number of genes not shared between the pair. Differences in gene content accumulate very rapidly at small core-genome distances but slow down at larger distances. At a core-genome diversity of 10^−3^, strains typically differ in a few hundred genes. These relationships are surprisingly

**FIG. 6.**
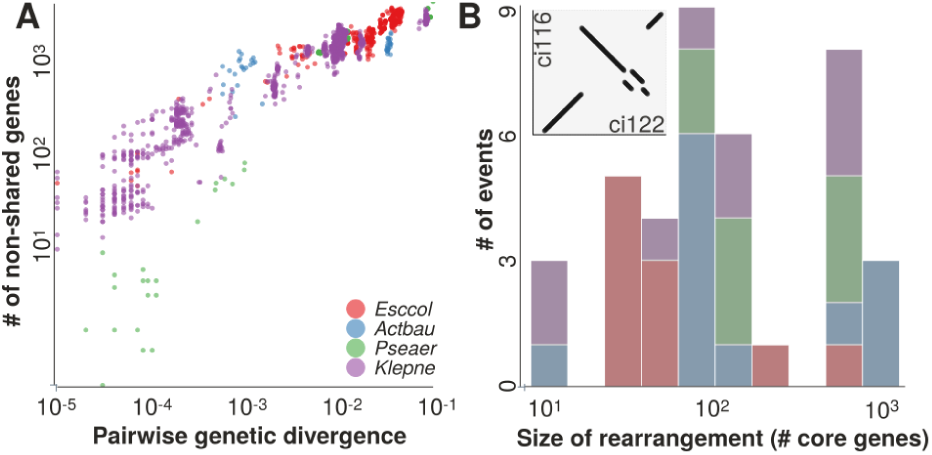
Gene gain/loss and genome rearrangements. A) Number of genes that aren’t shared between any pair of individuals in our sample scales sublinearly with the genetic distance of the corresponding core genome alignment. B) Histogram displaying the size distribution, in number of core genes, we detected in each isolate, relative to the most prevalent core gene order found for each species. Observed to be trimodal. Inset shows an example of a piece of an obtained dot plot for isolate 116 compared against isolate 122, both *A. baumannii.* This particular example was tabulated as 5 events.

#### consistent between different bacterial species

As we resolved the structure of each isolate genome, we also quantified the rearrangements with respect to the order of core genes for each isolate. The resulting histogram of the size of detected events is shown in Fig. 6B. Events were detected via breaks in the core genome order between isolates, see the inset of Fig. 6B for an example. In total we found 36 events. All but three events mapped to terminal branches on the core-genome tree. The remaining three being shared by two isolates each. This suggests that most rearrangements were recent and are pruned by purifying selection to maintain gene order. As most events were unique, we observed low correlation between genetic distance and number of rearrangements.

#### Ancestral recombination present in the core genome of *K. pneumoniae* and *E. coli*

Figure 7A,B display the core genome trees obtained for the *K. pneumoniae* and *E. coli* isolates from this study. Each tip is labeled with the associated MLST designations along with a colored circular marker. The estimated MLSTs are largely consistent with the clades defined by the tree. Additionally, we see strong correlations between the resistome of each isolate, shown to the right of the core tree, and its designated ST. However, the correspondence between resistome and the tree if far from perfect and we show below that the core-genome tree is inconsistent with large fractions of the core-genome diversity for both *K. pneumoniae* and *E. coli.*

**FIG. 7.**
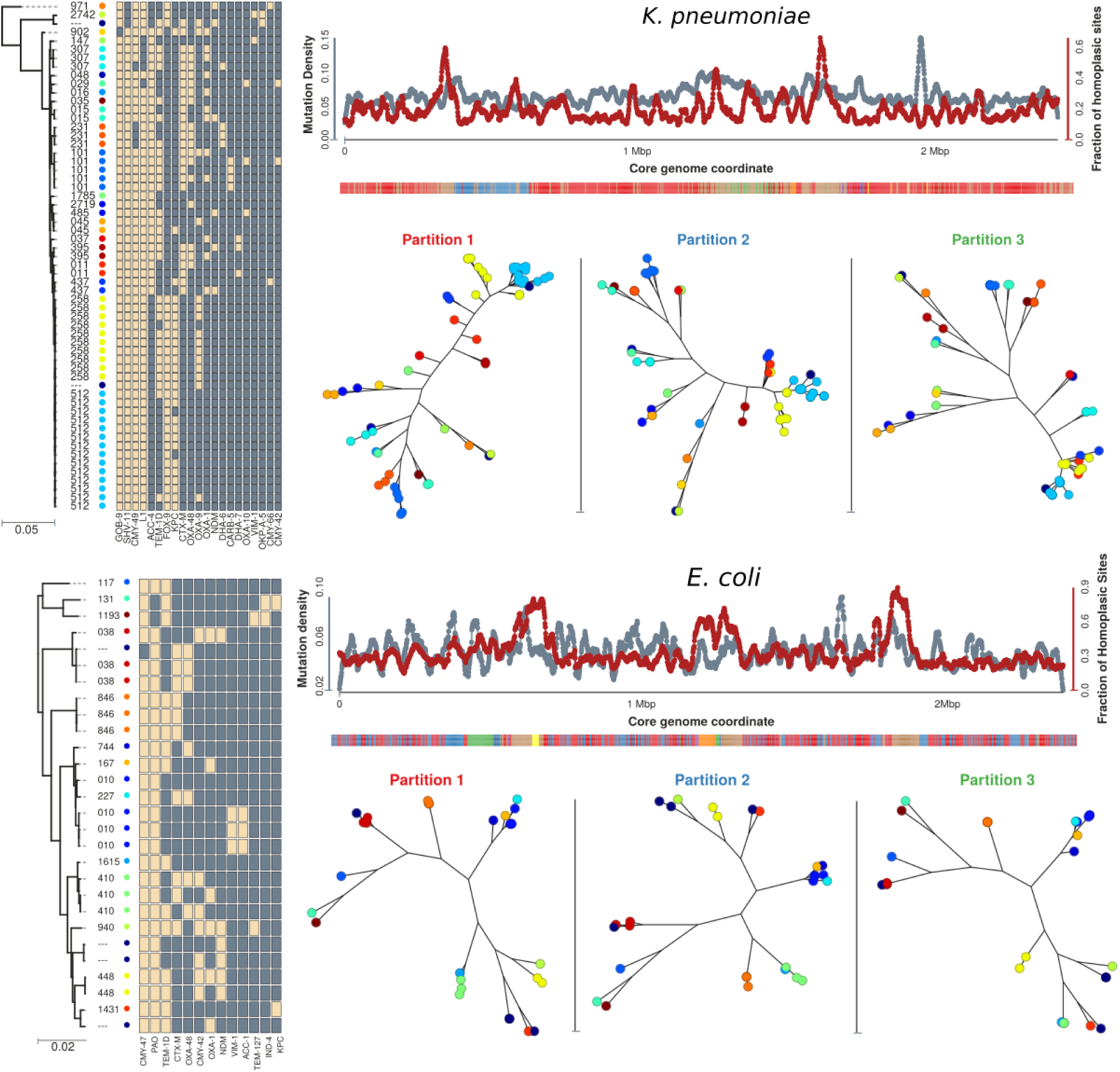
*K. pneumoniae* and *E. coli* are mosaics of ancestral recombination events. Core genome tree reconstruction from the alignment of core genes for *K. pneumoniae* (top) and *E. coli* (bottom). Each tip is labeled by its MLST and is given a corresponding marker for visual reference. Gene presence absence for carbapenemases, ESBLs, and other beta-lactams are plotted as a binary matrix to the right. Strong association between MLST and resistome. Density of polymorphic positions, as well as the fraction of such positions that were determined to be homoplasic, were computed along the core genome coordinate. Contiguous regions of high homoplasic density found for both *K. pneumoniae* and *E. coli.* This was corroborated by finding clusters of trees build from 5kb blocks with similar topology. Trees associated with the three most dominant partitions are shown. We note that the regions of anomolously large homoplasic fractions had a large variance in trees (shown as the tan color) and are likely homologous recombination hotspots.

To assess the degree to which genetic variation in the core-genome is compatible with a single vertical history as defined by the core-genome tree, we first looked for regions of high homoplasic density. Specifically, the ancestral state of each polymorphic position of the concatenated core genes was reconstructed for each internal node of the core-genome tree, using TreeTime (Sagulenko *et al.,* 2018). The number of mutations for each polymorphic site that were parsimoniously inferred on branches of the tree were then counted. If the evolutionary history at a given position is consistent with the single core genome tree, under an infinite sites model we expect to find each mutation exactly once in the tree; a reasonable approximation as the median distance between any two strains is small such that the probability that a position mutated independently on the tree is ≪ 1. Thus, if the position underwent recombination and hence has an evolutionary history that deviates from the consensus tree, this can manifest as a homoplasy - i.e. a mutation that appears on multiple branches within the tree. We note by itself, this is not a necessary and sufficient test for homologous recombination: an additional plausible hypothesis is simple convergent evolution due to the shared selection pressure applied on the clinical ecosystem.

Figure 7 shows the density of overall SNPs, along with the homoplasic density along the core-genome for *K. pneumoniae* and *E. coli,* chosen in the order previously discussed. In our sample of *K. pneumoniae* genomes, we found a basal fraction of a ~15% homoplasies relative to overall SNP density within the majority of the core genome. Additionally, we find two contiguous regions that have marked elevated rate of homoplasies suggesting either a large scale ancestral recombination event or a huge recombination hotspot. We note that one of these regions agrees well previous observations of that ST 258 is a recombinant mosaic of ST11 and ST42 (Chen *et al.,* 2014). In our sample of *E. coli* genomes, we find homoplasies are significantly more prevalent; we estimate ~ 30% of SNPs are inconsistent with the reconstructed core genome phylogeny, indicative of a higher homologous recombination rate. The fraction of homoplasies depends, however, on sample size and diversity such that these numbers are not directly comparable. Similar to our measurements of *K. pneumoniae,* we find three contiguous regions of high homoplasy density, likely to be ancestral recombination events, however the landscape is markedly more rugged in comparison.

To corroborate this result, we built local phylogenetic trees in 5 *kB* segments from the concatenated core genome in regions of collinearity, taking care not to extend a block across a rearrangement event as shown in the inset of Fig. 6, using FastTree (Price *et al.,* 2010). We performed an all-to-all comparison of the topology of each tree using the Robinson-Foulds metric. The resulting distance matrix of all pairwise comparisons was clustered into the dominant topologies, shown as different colors within the bar beneath the homoplasic density plot within Fig. 7 for *K. pneumoniae* and *E. coli.* Both assays are in good agreement with each other; wherever homoplasy density was anomalously high, we also resolved a substantially different tree relative to the background.

For clarity, we display the unrooted phylogenetic trees obtained from the tree largest partitions defined by the above analysis in Fig. 7 for *K. pneumoniae* and *E. coli.* Branch lengths are shown in log scale to better resolve intratype structure. The resulting trees immediately indicate the putative ancestral recombination events for each associated core genome partition. We first discuss the results obtained for *K. pneumoniae.* Partition two, shown in blue, represents a ~ 250*kB* region that differs from the global tree by two main topological changes: ST11 and ST437 both root within a subclade of ST258, and the clade containing ST971 and ST2742 directly attaches just above ST11. To our knowledge, this event has not been described in the literature. Partition three, shown in green, recapitulates the ancestral recombination event noted in (Chen *et al.,* 2014); we see ST15 now roots within ST258. Last we discuss the results obtained for *E. coli.* Specifically, partition two for *E. coli* differs from the background tree in that the clade containing ST410 and ST1615 now roots with ST846. Partition three of *E. coli* shows a transposition between ST846 and the clade containing ST131 and ST1193.

Taken together, ~ 30% and ~ 50% of the genomes of our sample of *K. pneumoniae* and *E. coli* respectively appears to be inconsistent with the core-genome consensus tree. However, at the present our analysis mostly was dominated by the signal of a handful of recombination events that presumably occurred before the carbapenemase became frequent. As such, the trees presented in Fig. 7 exhibit no significant intra-ST mixing that would be characteristic of homologous recombination.

Importantly we note that fraction of homoplasic SNPs is not well correlated with SNP density, exemplified best within the *K. pneumoniae* genome shown in Fig. 7. Transitions in tree topology are better predicted by sudden increases in homoplasy density than overall SNP density. Popular tools such as Gubbins (Croucher *et al.,* 2015) use stretches of high diversity to identify horizontally transfer events. This strategy likely works for transfer from distantly related strains, while common recombination between similar strains remain undetected. While the latter has a smaller impact on the reconstructed phylogeny, it might still mean that the history of any given locus is poorly approximated by the consensus tree. More sensitive methods will be necessary to detect and reconstruct the history of such loci. Similar analyses for *A. baumannii* and *P. aeruginosa* are presented in the S.I.

## DISCUSSION

Acquired antibiotic resistance of bacteria spreads via a complicated mix of processes involving (i) the clonal dissemination of pathogens, (ii) transfer of plasmids or homologous recombination, and (iii) mobilitization of the resistance gene to novel genomic backgrounds and non-homologous recombination (Sheppard *et al.,* 2016). The importance of these processes differ by resistance gene and species, but little is known about their quantitative relative importance.

Here, we have used long read sequencing to resolve the genomic background of a large number of carbapenemases genes isolated in the past decade in the University Hospital Basel. Our study substantially increases the number of publicly available high-quality complete genomes of carbapenemase-producing *enterobacteriaceae.* Such fully assembled genomes are necessary to develop quantitative surveillance methods to accurately measure, control, and ultimately predict the global spread of bacterial resistance.

A crucial question that arises when studying the epidemiology of carbapenemase genes is what relevant genomic unit should be tracked. Carbapenemase genes themselves lack resolution, while MLST types don’t track the genes due to frequent gene mobilization and horizontal transfer. Our results suggest that the structure of plasmids or the chromosomal neighborhood of carbapenemase genes can be efficiently used to reconstruct their relationships and is informative on the relevant time scales. Such inference based on genome structure/syntheny alignments can be performed efficiently and could be supplemented by nucleotide information if necessary.

High quality bacterial genomes currently require hybrid assembly of accurate short sequencing reads and long-reads (either from ONT or PacBio technologies) for long range order (Loman and Pallen, 2015). Short-read only assemblies won’t resolve the genomic context of carbapenemase genes beyond a few genes due to the repetitive nature of their neighborhoods (compare Fig. 3). Long-read only assemblies, by contrast suffer from indel errors. Recent improvements in long-read sequencing technologies have made large scale assembly of complete genomes possible in a cost-effective manner. We routinely achieved sufficient coverage to assemble 12 genomes on a single ONT flow cell and future improvements will surely reduce the price even more. After a decade during which almost all newly sequenced bacterial genomes were fragmented short read assemblies, we expect the availability and diversity of fully assembled genome to improve dramatically. A comprehensive analysis of the complex interplay of horizontal transfer, genome rearrangements, recombination, and sequence evolution by mutation will require new models and theoretical frameworks. We need computationally efficient tools to analyze large, diverse collections of whole genomes and extract quantitative determinants of the spread of resistance genes and plasmids. This remains an important challenge for future research.

## ACKNOWLEDGEMENT

We are grateful to Eduardo Rocha and Zamin Iqbal for stimulating discussions and constructive comments on the manuscript. Computational work was performed at sciCORE scientific computing center at University of Basel. The study was funded by the University of Basel.

